# Start-Stop Assembly: a functionally scarless DNA assembly system optimised for metabolic engineering

**DOI:** 10.1101/361626

**Authors:** George M. Taylor, Paweł M. Mordaka, John T. Heap

## Abstract

DNA assembly allows individual DNA constructs or designed mixtures to be assembled quickly and reliably. Most methods are either: (i) Modular, easily scalable and suitable for combinatorial assembly, but leave undesirable ‘scar’ sequences; or (ii) bespoke (non-modular), scarless but less suitable for construction of combinatorial libraries. Both have limitations for metabolic engineering. To overcome this trade-off we devised Start-Stop Assembly, a multi-part, modular DNA assembly method which is both functionally scarless and suitable for combinatorial assembly. Crucially, 3 bp overhangs corresponding to start and stop codons are used to assemble coding sequences into expression units, avoiding scars at sensitive coding sequence boundaries. Building on this concept, a complete DNA assembly framework was designed and implemented, allowing assembly of up to 15 genes from up to 60 parts (or mixtures); monocistronic, operon-based or hybrid configurations; and a new streamlined assembly hierarchy minimising the number of vectors. Only one destination vector is required per organism, reflecting our optimisation of the system for metabolic engineering in diverse organisms. Metabolic engineering using Start-Stop Assembly was demonstrated by combinatorial assembly of carotenoid pathways in *E. coli* resulting in a wide range of carotenoid production and colony size phenotypes indicating the intended exploration of design space.

**GRAPHICAL ABSTRACT:** 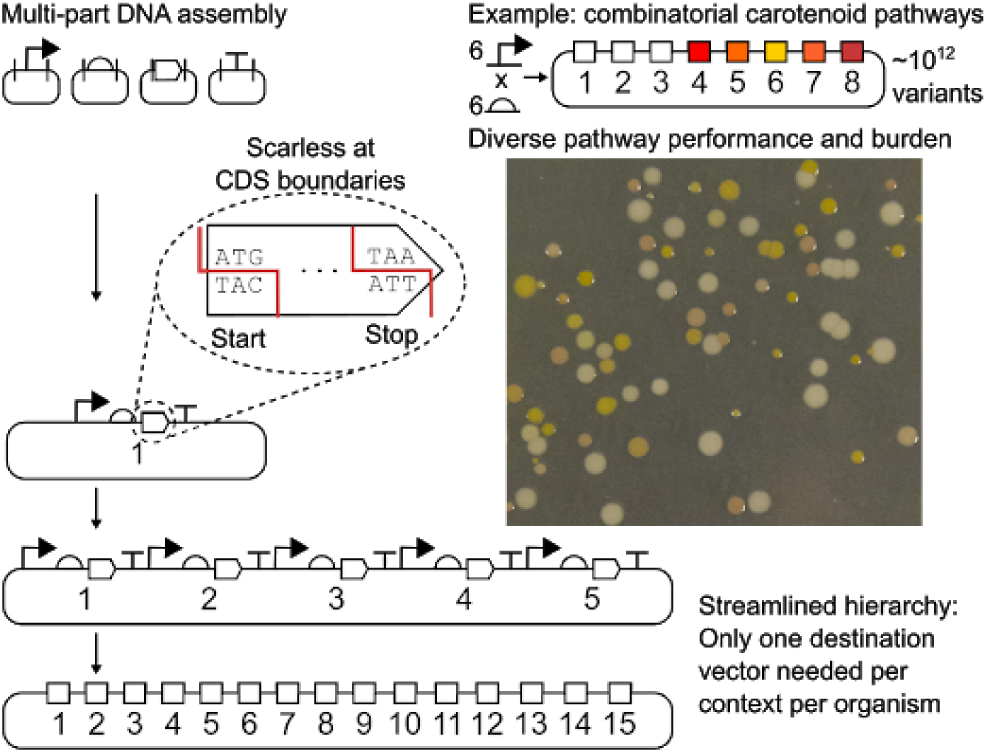

## INTRODUCTION

In recent years, new types of DNA cloning methods often referred to as DNA assembly have been reported which are much more suitable for combining multiple DNA sequence elements or ‘parts’ in a single step than conventional restriction-ligation cloning (1). These multi-part DNA assembly methods can mostly be categorised into one of two types: (i) Modular approaches, in which a framework of specified steps, rules and design constraints including pre-defined module formats allows highly efficient multi-part assembly of individual constructs or designed combinatorial mixtures of constructs. However, these benefits come at the cost of fixed ‘scar’ sequences at junctions between parts in the assembled construct and a substantial upfront investment of effort and resources to obtain a required set of suitable assembly vectors, and to prepare sequences of interest in the appropriate format. (ii) Bespoke approaches, which are flexible, require little upfront effort to establish or planning of future steps, and typically involve an *ad hoc* approach to the design of cloning strategies which is intuitively more similar to conventional cloning approaches. However, these non-modular approaches depend on sequence overlaps to direct assembly, so they have a greater need for custom oligonucleotide primers (typically one pair per junction between parts) and sequence verification (due to PCR steps) and are less suitable for combinatorial assembly due to lower efficiency (meaning fewer clones are obtained following transformation so smaller libraries are generated) and greater potential for bias particularly due to repetitive sequences. Modular multi-part DNA assembly methods include Golden Gate assembly (2–4) (and variants (4–14)), BASIC (15), BioBrick assembly (16) (and variants such as BglBrick (17, 18)) and Gateway cloning (19–21). Bespoke multi-part DNA assembly methods include Gibson Assembly (22), AQUA cloning (23), Twin Primer Assembly (24), ligase cycling reaction (25), SLIC (26), SLiCE (27), overlap extension PCR (28) and CPEC (29). Multi-part DNA assembly has quickly become important in synthetic biology, enabling an increase in the scale, scope and speed of studies (1).

The development of effective metabolic pathway-encoding constructs is a key application for multi-part DNA assembly. The amount of a protein produced by an expression construct depends on the expression control parts used (particularly promoters, ribosome-binding sites (RBSs) and transcriptional terminators) as well as the coding sequences (CDSs), which affect expression through mRNA structure and codon usage. The design of a multi-protein expression construct, such as those encoding metabolic pathways, involves a multi-dimensional ‘design space’, in which the concentration of each protein represents a dimension (30–34). Within any such design space there will be regions of functional pathway-encoding designs, including local optima, but also regions of poorly-functional or non-functional pathway-encoding designs. Natural metabolic pathways, and the sequences encoding them, are optimised by natural selection for efficient formation of the intended product, and to avoid deleterious effects on the cell (35). In contrast, heterologous or synthetic metabolic pathways can be non-optimal in various ways including poor flux through the pathway to the intended product, the formation of by-products, the accumulation of intermediates (which may be toxic), excessive enzyme overexpression (36, 37) and deleterious effects on the cell caused by one or more of these issues (32, 34, 36, 38, 39). These issues depend largely on the profile of absolute and relative concentrations of the enzymes in a pathway. In principle, metabolic modelling methods allow prediction of suitable enzyme concentration profiles for a particular pathway in a given host organism. In practice, predictive modelling approaches are generally demanding, requiring suitable metabolic models and parameter values which may not be available. Furthermore, even if an optimal enzyme concentration profile is known, at present it is difficult or impossible to rationally design an individual DNA sequence to express a protein of interest to a specific target concentration (30), a problem which increases in difficulty with increasing numbers of proteins in a multi-protein expression construct (30, 31).

Modular multi-part DNA assembly facilitates generally-applicable strategies to identify and optimise metabolic pathway-encoding constructs from among the many possible designs for any given pathway. Firstly, efficient assembly is not limited to construction of individual expression constructs, but can be used to generate combinatorial libraries of multi-protein expression constructs, in which the expression of each protein is varied independently, or partially-independently, by using mixtures of expression control parts instead of individual expression control parts (31, 32). The design of such mixtures (in terms of the number of parts and the distribution of their expression levels) will determine the distribution and granularity of sampling of design space with respect to the relevant protein, allowing for tunable search strategies effective for large design spaces, limited primarily by the throughput of screening once host cells are transformed with the library. Secondly, once functional metabolic pathway-encoding construct designs are identified, modular DNA assembly facilitates rational optimisation of these through further, more fine-grained rounds of combinatorial searches, and/or by manually-designed substitutions of specific expression control parts (31). Relative to predictive modelling approaches, these approaches are pragmatic and empirical, requiring little a *priori* knowledge of suitable enzyme concentrations, nor of the protein concentrations that will result from particular combinations of expression control parts and CDSs. The main requirements for these approaches are suitable expression control parts which span an appropriate range of strengths, and a suitable modular multi-part DNA assembly method.

Of the modular DNA assembly methods, Golden Gate assembly (2, 3) and variants (4–14) are particularly widely used, including for construction of metabolic pathway-encoding constructs (10, 14, 40–42). Golden Gate uses type IIS restriction endonucleases, which are similar to classical type II restriction endonucleases, except their restriction site is offset from their recognition site, rather than within it. IIS restriction endonuclease sites are asymmetrical, which confers directionality. Type IIS restriction endonucleases, in combination with appropriately-designed DNA sequences, can be used to generate DNA fragments with any desired cohesive end sequences (of the 4^*n*^ possibilities, where 4 reflects the four bases A, T, C or G; and *n* is the length of the overhang) independently of the sequence of the recognition site. The cohesive end sequences can be non-palindromic, avoiding the self-ligation of palindromic cohesive ends like those generated by classical type II endonucleases. Golden Gate assembly uses a series of DNA fragments generated in this way by a single type IIS endonuclease (in each assembly step), such that each fragment has unique non-palindromic cohesive ends designed to anneal only to the correct end of the next DNA fragment in the series. Such a series of DNA fragments can be efficiently and directionally assembled, in the intended order, by ligation. Furthermore, an efficient ‘one-pot’ assembly reaction containing both ligase and the appropriate type IIS endonuclease can be used (2, 3), because unique non-palindromic cohesive ends can only either (i) ligate to the intended cognate cohesive DNA end, in which case the restriction site is not regenerated, so the ligation product is not subsequently restricted; or (ii) re-ligate to the donor plasmid fragment from which they were originally excised, in which case the site is regenerated and can be subsequently restricted again. The overall effect is that the concentration of the intended assembly product in the one-pot reaction increases over time, whereas the concentration of original or re-ligated donor plasmids decreases over time, resulting in very efficient assembly (2, 3). One-pot assembly reactions can be performed under isothermal conditions, but are more efficient if the reaction temperature is cycled between the different optimal temperatures of the ligase and endonuclease (3). The Golden Gate approach as originally described (2) effectively defines a broad strategy, rather than specific details of how it might generally be applied. Subsequent publications from various groups have described particular implementations of Golden Gate, crucially including frameworks for hierarchical assembly of numerous parts, and collections of vectors and parts in the relevant formats for these (4, 5). These implementations are mostly focused on specific organisms, such as plants (4–7), yeast (8–12) or *Escherichia coli* (13, 14). There is a lack of modular, multi-part assembly systems which are widely-applicable to many organisms with a minimum of cloning effort, which would be very useful.

All the above Golden Gate approaches require specific ‘fusion site’ sequences to be defined for the unique cohesive ends, which are incorporated into the assembled construct as unavoidable scar sequences at the junctions between parts. Scars therefore impose design constraints on constructs assembled by Golden Gate and other modular multi-part DNA assembly methods. At the junction between some types of parts, scars can have a substantial impact on functional properties. A key example is the junction between a CDS and the upstream sequence. Scars at this junction are transcribed to mRNA, potentially affecting mRNA structure (which in turn affects functional properties (43)), RBS accessibility, and may be an issue for regulatory approaches achieved using designed RNA elements, such as riboswitches (44), riboregulators (45) and small transcription activating RNAs (STARs) (46). Scars at this junction are also within the region of mRNA bound by the ribosome, so also directly affect ribosome binding and therefore affect initiation of translation through this mechanism. To illustrate this, we performed a direct comparison of the impacts of several assembly scars at a junction between a CDS and the upstream sequence (Figure S1).

It is preferable to minimise constraints on the design of DNA constructs generally, but particularly in cases where the desired functional properties of the construct are likely to be sensitive to the absolute and/or relative expression levels of its CDSs, and therefore sensitive to the presence of scars. An important example is the assembly of individual or combinatorial mixtures of metabolic pathway-encoding constructs. A DNA assembly system providing efficient modular multi-part assembly, yet without leading to scars at sensitive positions, would be widely useful. Here we develop such a DNA assembly system, called Start-Stop Assembly, and demonstrate its utility for combinatorial assembly of metabolic pathway-encoding constructs.

## MATERIALS AND METHODS

Additional materials and methods are available in the Supplementary Materials.

### Bacterial strains and growth conditions

*E. coli* strain DH10B was used for all DNA assembly, all other cloning, and all other experiments described in this study. *E. coli* DH10B cells were routinely grown in LB medium (tryptone 10 g l^−1^, yeast extract 5 g l^−1^ and NaCl 5 g l^−1^) at 37 °C, with shaking at 225 r.p.m, or on LB agar plates (containing 15 g l^−1^ bacteriological agar) at 37 °C. LB was supplemented with ampicillin (100 μg ml^−1^), tetracycline (10 μg ml^−1^), kanamycin (50 μg ml^−1^) or chloramphenicol (25 μg ml^−1^) as appropriate. *E. coli* cells were routinely transformed by electroporation (47).

### Plasmid construction

Except where stated otherwise, plasmid construction was carried out using standard molecular cloning methods. Details are provided in the Supplementary Materials, tables of plasmids (Table 1 and Table S3), oligonucleotides (Table S4) and synthetic DNA (Table S6). This includes a suggested method to construct alternative Level 2 destination vectors (Figure S6).

**Table 1.** List of core Start-Stop Assembly vectors. The 15 vector names reflect their level in the hierarchy (number in blue) and their donor fusion sites (letters or numbers in red) or in the case of Level 3 vectors their acceptor fusion sites (numbers in green). Each vector contains an assembly cassette (including a *lacZα* gene for blue/white screening), a resistance marker (AmpR, TetR, KanR or CamR) and a replicon (high-copy pMB1 or low-copy p15A). Each also has a unique ID number (pGT…). Genbank accession numbers are shown.

| Plasmid Name | Accession Number | Level | Acceptor fusion sites | Donor fusion sites | Comments |
| --- | --- | --- | --- | --- | --- |
| pStA**0** | MG649420 | **0** | F-R (Bsal) | α-ε (SapI) | AmpR, pMB1, *lacZα*, ID = pGT400 |
| pStA**1**AZ | MG649422 | **1** | α-ε (SapI) | **A-Z** (Bsal) | TetR, p15A, *lacZα*, ID = pGT401 |
| pStA**1**AB | MG649421 | **1** | α-ε (SapI) | **A-B** (Bsal) | TetR, p15A, *lacZα*, ID = pGT402 |
| pStA**1**BZ | MG649424 | **1** | α-ε (SapI) | **B-Z** (Bsal) | TetR, p15A, *lacZα*, ID = pGT403 |
| pStA**1**BC | MG649423 | **1** | α-ε (SapI) | **B-C** (Bsal) | TetR, p15A, *lacZα*, ID = pGT404 |
| pStA**1**CZ | MG649426 | **1** | α-ε (SapI) | **C-Z** (Bsal) | TetR, p15A, *lacZα*, ID = pGT405 |
| pStA**1**CD | MG649425 | **1** | α-ε (SapI) | **C-D** (Bsal) | TetR, p15A, *lacZα*, ID = pGT406 |
| pStA**1**DZ | MG649428 | **1** | α-ε (SapI) | **D-Z** (Bsal) | TetR, p15A, *lacZα*, ID = pGT407 |
| pStA**1**DE | MG649427 | **1** | α-ε (SapI) | **D-E** (Bsal) | TetR, p15A, *lacZα*, ID = pGT408 |
| pStA**1**EZ | MG649429 | **1** | α-ε (SapI) | **E-Z** (Bsal) | TetR, p15A, *lacZα*, ID = pGT409 |
| pStA**2**12 | MG649430 | **2** | A-Z (Bsal) | **1-2** (BbsI) | KanR, p15A, *lacZα*, ID = pGT417 |
| pStA**2**23 | MG649431 | **2** | A-Z (Bsal) | **2-3** (BbsI) | KanR, p15A, *lacZα*, ID = pGT418 |
| pStA**2**34 | MG649432 | **2** | A-Z (Bsal) | **3-4** (BbsI) | KanR, p15A, *lacZα*, ID = pGT419 |
| pStA**3**13 | MG649433 | **3** | **1-3** (BbsI) | N/A | CamR, p15A, *lacZα*, ID = pGT415 |
| pStA**3**14 | MG649434 | **3** | **1-4** (BbsI) | N/A | CamR, p15A, *lacZα*, ID = pGT416 |

### Storing genetic parts

Genetic parts were cloned in Level 0 vector pStA0 using one-pot restriction-ligation assembly reactions (as described below) of inserts obtained by PCR amplification or DNA synthesis, or by inverse PCR using pStA0 as template and primers incorporating the part sequence. Spacer parts were generated as double-stranded oligonucleotide linkers by oligo annealing, and were not cloned for storage. See Supplementary Materials for details.

### Start-Stop Assembly reactions

Start-Stop Assembly reactions contained 20 fmol of destination vector plasmid DNA, 40 fmol of each insert (plasmid DNA or annealed oligonucleotides), T4 DNA Ligase buffer, 400 units of T4 DNA Ligase and 10 units of the appropriate restriction endonuclease (SapI, BsaI or BbsI) in a total reaction volume of 20 μl. All enzymes and T4 DNA Ligase buffer were supplied by New England Biolabs. Reactions were incubated using a thermocycler for 30 two-step cycles of 37 °C for 5 minutes then 16 °C for 5 minutes, before a single final denaturation step at 65 °C for 20 minutes.

### Analysis of Level 1 assembly fidelity and bias by flow cytometry

*E. coli* DH10B cells were transformed with Start-Stop Assembly reactions. Isolated transformant clones or pools of transformant clones were used to inoculate 5 ml LB broth supplemented with the appropriate antibiotic, which was cultured overnight (16 h) at 37 °C with shaking at 225 r.p.m. For flow cytometry analysis, overnight cultures were subcultured by 1:1000 dilution into 5 ml fresh LB medium with the appropriate antibiotic and grown for 6 h at 37 °C with shaking at 225 r.p.m. Cultures were diluted 1:50 in filtered PBS and immediately subjected to flow cytometer analysis.

Fluorescence was measured using an Attune NxT flow cytometer (Invitrogen). The voltage gains for each detector were set to ensure all constructs fell within the dynamic range: FSC, 440 V; SSC, 440 V; BL1, 500 V. 10,000 events gated by forward scatter height (FSC-H) and side scatter height (SSC-H) that represent the *E. coli* cell population were collected at 100 μl min^−1^ sample flow rate. Flow cytometry data were analysed using FlowJo (http://flowjo.com). Cells were gated by FSC-H and SSC-H and the geometric mean from the BL1 detector was exported as the fluorescence value.

### Coding sequences for carotenoid metabolic pathway libraries

The eight CDSs used in the assembly of carotenoid metabolic pathway libraries were *dxs* (encoding 1-deoxyxylulose-5-phosphate synthase; WP_074468184) from *E. coli* MG1655, *crtE* (encoding geranylgeranyl pyrophosphate synthase; WP_010888034) from *Deinococcus radiodurans* R1, *crtB* (encoding phytoene synthase; WP_010887508) from D. *radiodurans* R1, *crtI* (encoding phytoene desaturase; WP_010887507) from *D. radiodurans* R1, *idi* (encoding isopentenyl pyrophosphate isomerase; AAC32208) from *Haematococcus pluvialis*, *lcyB* (encoding beta-lycopene cyclase; ABR57232) from *Solanum lycopersicum*, *crtZ* (encoding β-carotene hydroxylase; WP_072137426) from *Pantoea ananatis* and *crtW* (encoding β-carotene ketolase; BAB74888) from *Nostoc sphaeroides* PCC 7120. Each CDS was codon-optimised for expression in *E. coli* using the OPTIMIZER guided random methodology (48). Iterations of the optimisation were repeated until each CDS had a codon adaptation index (CAI) score >0.65. Each CDS was synthesised as a linear gBlock DNA fragment (Integrated DNA Technologies).

### Analysis of phenotypic diversity of carotenoid pathway library clones by image analysis

*E. coli* DH10B cells were transformed with 5 μl of Start-Stop Assembly reaction product, allowed to recover for one hour, plated and grown at 30 °C for 48 h. Transformation plates were photographed under consistent conditions using a Canon 60D camera. The images were analysed using OpenCFU v.3.8 beta (49), and colonies were detected using an inverted threshold of 3 and a minimum radius of 10 pixels and a maximum radius of 500 pixels. The area and red, green, blue colours of each colony were determined.

### Quantification of lycopene content of cells

*E. coli* was cultured in tubes containing 5 ml LB supplemented with the appropriate antibiotics and grown at 37 °C at 225 r.p.m for 24 h. To obtain the cry cell weight (DCW), after 24 h cultivation 2 ml of liquid culture was centrifuged at 14,000 rpm for 5 minutes to remove supernatant, the bacterial cell pellet was washed in ddH_2_O and then centrifuged again, and dried at 70 °C until a constant weight was obtained. To obtain lycopene concentrations, following 24 hours incubation 2 ml of liquid culture was centrifuged at 14,000 rpm for 5 minutes to remove supernatant, the bacterial cell pellet was washed in ddH_2_O and then centrifuged as before. The bacterial cell pellet was resuspended in 1 ml acetone and incubated at 55 °C for 15 minutes to extract lycopene. The supernatant was obtained by filtration through a 0.22 μm pore-size nylon membrane for LC-DAD analysis.

Lycopene was detected and measured using an Agilent LC system with UV/Vis diode array detector. Absorbance at 450 nm and 471 nm were monitored and the peak area corresponding to each component integrated to provide a measure of abundance. The LC column used was an Acquity UPLC Peptide BEH C18 column (2.1 × 100mm, 1.7 μm, 300Å, Waters). LC buffers were 50% of methanol in water (A) and 25% of ethyl acetate in acetonitrile (B). All the solvents used were HPLC grade. The LC method was 6.5 minutes in total with 1.5 minutes of post run time (0-1 min: 30% A, 70% B; 1-6 min: 0.1% A, 99.9% B; 6-6.5 min: 30% A, 70% B; at a flow rate of 0.3 ml/min). The injection volume for the samples was 1 μl. Commercially available lycopene (Sigma-Aldrich) was dissolved in acetone as a standard and a standard curve was generated.

## RESULTS

### A strategy for functionally-scarless assembly of expression units

Normally, modular multi-part DNA assembly methods necessarily generate scars, because sequence identity between DNA ends is required to direct recognition and joining of the correct DNA ends, so these fusion site sequences become incorporated into the assembled construct. We devised a strategy to mitigate this issue. We reasoned that if existing constraints on design of expression constructs included conserved sequence motifs which could also be used as fusion sites, then DNA assembly using these fusion sites would be effectively scarless at the relevant part junctions, as no additional constraints would be introduced by assembly. Such motifs would need to be very well conserved, located at boundaries between functional elements (to be useful for assembly) and which are sensitive to scars (to benefit from a scarless design), and compatible with the length of overhang of a cohesive DNA end generated by a known type IIS restriction endonuclease. We identified start and stop codons as the key natural constraints that meet these requirements. Start and stop codons can be completely conserved and occur at the beginning and end of every CDS, sites which are both boundaries between functional elements and particularly sensitive to scars. The type IIS restriction endonucleases employed in modular DNA assembly methods are usually those which generate cohesive DNA ends with 4 bp overhangs, such as BsaI, BbsI or BsmBI (2, 50), whereas start and stop codons are 3 bp long. In order to use start and stop codons as fusion sites in scarless designs, a type IIS restriction endonuclease which generates 3 bp overhangs would be required. Several such endonucleases are known, including the commercially-available enzyme SapI, which we use here.

Starting with the core concept of start and stop codons as fusion sites, we designed a generic ‘expression unit’ architecture to serve as the first assembly level (Level 1) in a hierarchical, multi-part, modular DNA assembly framework (Figure 1a) called Start-Stop Assembly, which builds on the principles of Golden Gate assembly. Each expression unit for a single protein is divided into four parts; the promoter, UTR/RBS, CDS and transcriptional terminator; which are key functional modules to allow control of transcription and translation as independently as possible. These four modules are delineated from one another and from a vector by five fusion sites (Figure 1a) which we named α (alpha), β (beta), γ (gamma), δ (delta) and ε (epsilon). To allow multi-part assembly in a single one-pot restriction-ligation reaction using a single type IIS restriction enzyme, the fusion sites must all be of the same length. First we defined the γ fusion site as the sequence ATG because it is the most common (51, 52) and most efficient (53) start codon, and we defined the δ fusion site as the sequence TAA because it is the most frequently-used and efficient (54) stop codon. The β fusion site is the junction between the promoter and the UTR/RBS, where the transcriptional start site (TSS) is found. In an attempt to minimise any potential impact of the β fusion site as a scar in assembled constructs, we sought to identify a consensus sequence for the *E. coli* TSS which could be used to define the β fusion site. The sequences of 3746 TSSs previously described in *E. coli* MG1655 (55) were aligned (Figure S2). A bias in the frequency of bases was evident at two of the three positions, but no overall consensus was identified. Therefore the sequence CCA was assigned to the β fusion site, by considering both the frequencies of bases at each position in the TSS alignment and minimising similarity with the previously defined γ and δ fusion sites (Figure S2). There are no natural constraints to guide definition of the sequences of the α and ε fusion sites, which are the external junctions of each expression unit with the promoter or terminator, respectively, so we simply designed these to be as different as possible to the three fusion sites already defined (β, γ and δ) in order to minimise mis-assemblies. CAG was assigned to the α fusion site and GGA to the ε fusion site (Figure 1a).

**Figure 1.**
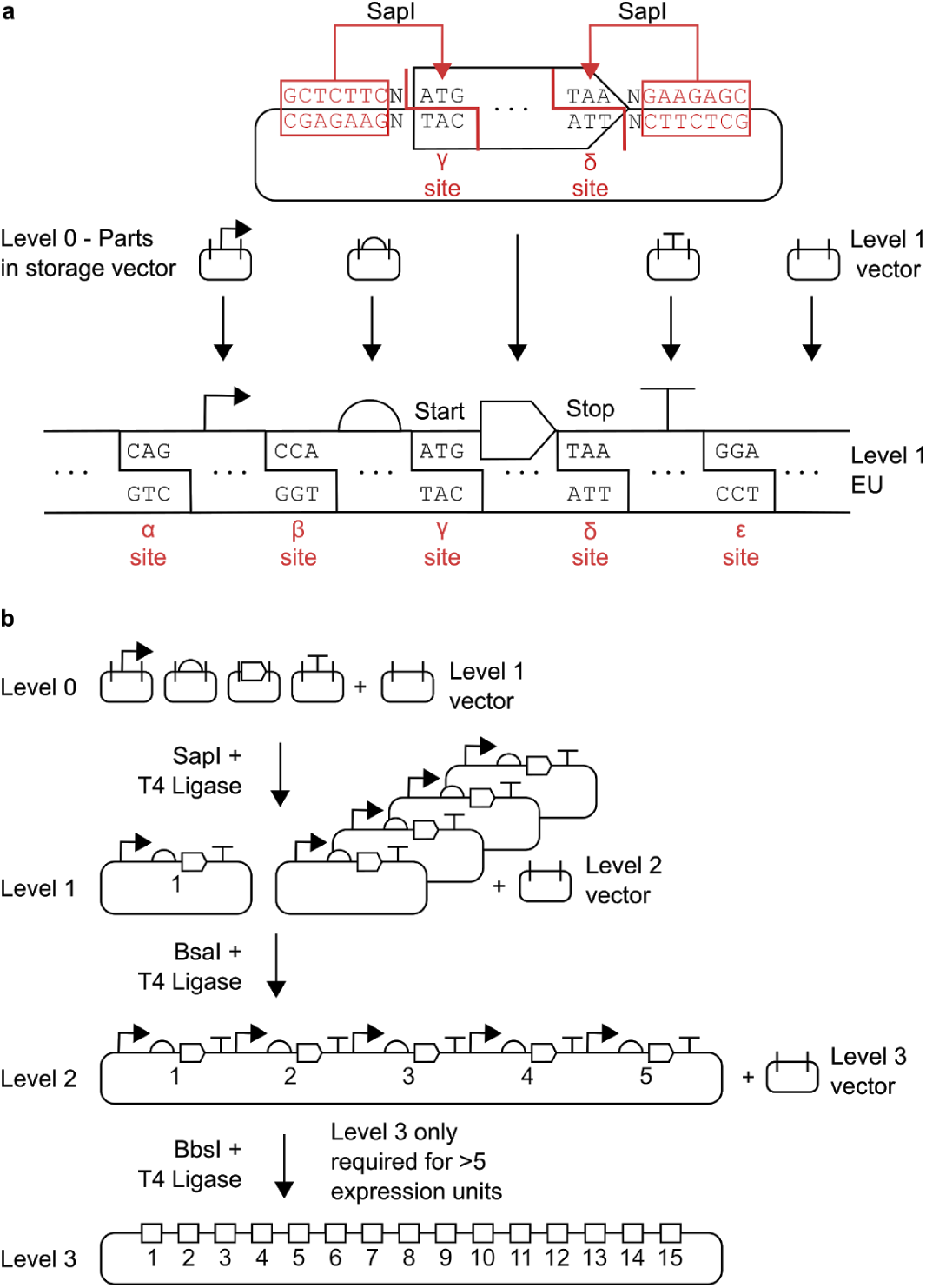
Schematic overview of Start-Stop Assembly. **(a)** Architecture of Level 1 assembly of expression units from genetic parts, showing detail for the CDS part in Level 0. Parts stored in Level 0 are flanked by inward-facing SapI restriction sites. Each type of part (promoter, RBS, CDS, terminator) uses different and unique fusion sites (α-ε sites) that allow the correct assembly of expression units. Coding sequences use the start codon ATG (γ site) and stop codon TAA (δ site) as fusion sites allowing functionally scarless assembly of expression units, **(b)** Schematic illustration showing the overall framework for Start-Stop Assembly. Up to 15 expression units can be hierarchically assembled from basic parts.

### Requirements for Level 1 assembly of expression units from Level 0 parts

To assemble expression units using the Level 1 architecture described above, and to ensure compatibility with efficient one-pot restriction-ligation assembly reactions, it is necessary for DNA parts (of each type) and vectors to meet certain specifications. Each DNA part must begin and end with the corresponding fusion sites for the type of part, and in the case of the β, γ and δ sites these should be positioned at the TSS (if known), start codon or stop codon, respectively. Each part must also be flanked by an inward-facing SapI recognition site (directionality exists because the recognition site GCTCTTC is asymmetrical) separated from the fusion site by a 1 bp spacer (Figure 1a, Figures S7-S10). This arrangement ensures that excision of each part by SapI will result in cohesive DNA ends with 3 bp 5’ overhangs corresponding to the 3 bp fusion sites. Appropriately-formatted DNA parts could be generated by any applicable method such as PCR or DNA synthesis, and could be used for assembly as either linear DNA or circular plasmid DNA molecules. If a part is cloned into a DNA vector, which is typical, the vector should not contain any other SapI sites in order to maximise the efficiency of one-pot restriction-ligation assembly reactions and avoid misassembly and bias. DNA parts should not contain any internal SapI sites, as these are used in Level 1 assembly; nor any BsaI sites or BbsI sites, as these are used in later assembly levels. These specifications define Level 0 (zero) of Start-Stop Assembly, which represents the initial ‘entry’ or ‘storage’ level of the system.

Instead of numerous entry or storage vectors as required by some systems, we designed and constructed a single Level 0 vector, named pStA0, for storage of parts of all types in *E. coli* (Table 1). For convenience, pStA0 includes a *lacZα* gene for blue/white screening and outward-facing BsaI restriction sites for efficient cloning of individual parts using special storage acceptor fusion sites (F and R; Figure S3, Note S2). To facilitate storage of parts in Level 0 in the required format, we have defined standard prefix and suffix sequences (Table S1) to be added to parts either as primer ‘tails’ (Table S2) when PCR-amplifying parts for storage, or when designing part sequences for synthesis. These sequences include inward-facing BsaI recognition sites and storage donor fusion sites for cloning formatted parts into Level 0 storage vector pStA0, as well as inward-facing SapI recognition sites with α, β, γ, δ or ε donor fusion sites (depending on the type of part) for subsequent multi-part assembly of expression units in Level 1 (Figure S7-10, Table S1). Finally, assembly requires a Level 1 vector including two outward-facing SapI recognition sites with appropriately-positioned α and ε acceptor fusion sites, but lacking any other SapI sites, for the reasons described above. These details are made clear by Figure S11.

### Validation of multi-part DNA assembly using 3 bp fusion sites

The performance of multi-part DNA assembly frameworks using IIS restriction endonucleases and typical 4 bp fusion sites is well validated in published studies (3, 4), whereas 3 bp fusion sites are rarely used. Our proposed Level 1 architecture for Start-Stop Assembly using five 3 bp fusion sites (Figure 1a) might cause performance differences to existing methods in terms of fidelity or bias of assembly. Therefore we set out to evaluate and validate assembly using this Level 1 design alone, independently of a multi-level hierarchical assembly. For this purpose, it was first necessary to obtain suitable genetic parts in the Level 0 format described above, and a Level 1 vector. A set of six promoters (Table S3) was chosen from a widely-used (56) library of constitutive *E. coli* promoters (57). These six promoters were chosen on the basis of previously-reported characterisation data (57) to span a wide range of expression strengths, and to be evenly spaced within that range (Figure S5a). Next, we constructed an RBS library by PCR using degenerate primers and characterised 96 individual RBSs (Figure S4). Six of these RBSs (Table S3) were chosen to give a wide and evenly-spaced distribution of expression strengths (Figure S4 and S5b). Four strong terminators (Table S3) with dissimilar sequences (to minimise the chance of undesirable homologous recombination) were chosen from a previously-characterised library of terminators (58). To facilitate simple visualisation and measurement of expression, a reporter gene encoding enhanced yellow fluorescent protein (EYFP) was chosen as a CDS. The chosen promoters, RBSs, terminators and EYFP-encoding CDS *eyfp* were cloned into pStA0 (as described in Supplementary Materials and Methods) ready for use in assembly. We designed and constructed several Level 1 vectors (Table 1) which are described in more detail later in this paper. For the purposes of these initial experiments to validate Level 1 assembly, the key features of Level 1 vectors are the two outward-facing SapI recognition sites with appropriately-positioned α and ε fusion sites (Figure S11) and a *lacZα* gene between the SapI sites to allow blue/white screening. Two types of assembly experiments were designed to validate Level 1 assembly, the first using replicates of assembly of an individual expression construct to assess fidelity (Figure 2a), and the second using a combinatorial assembly to assess bias (Figure 2b).

**Figure 2.**
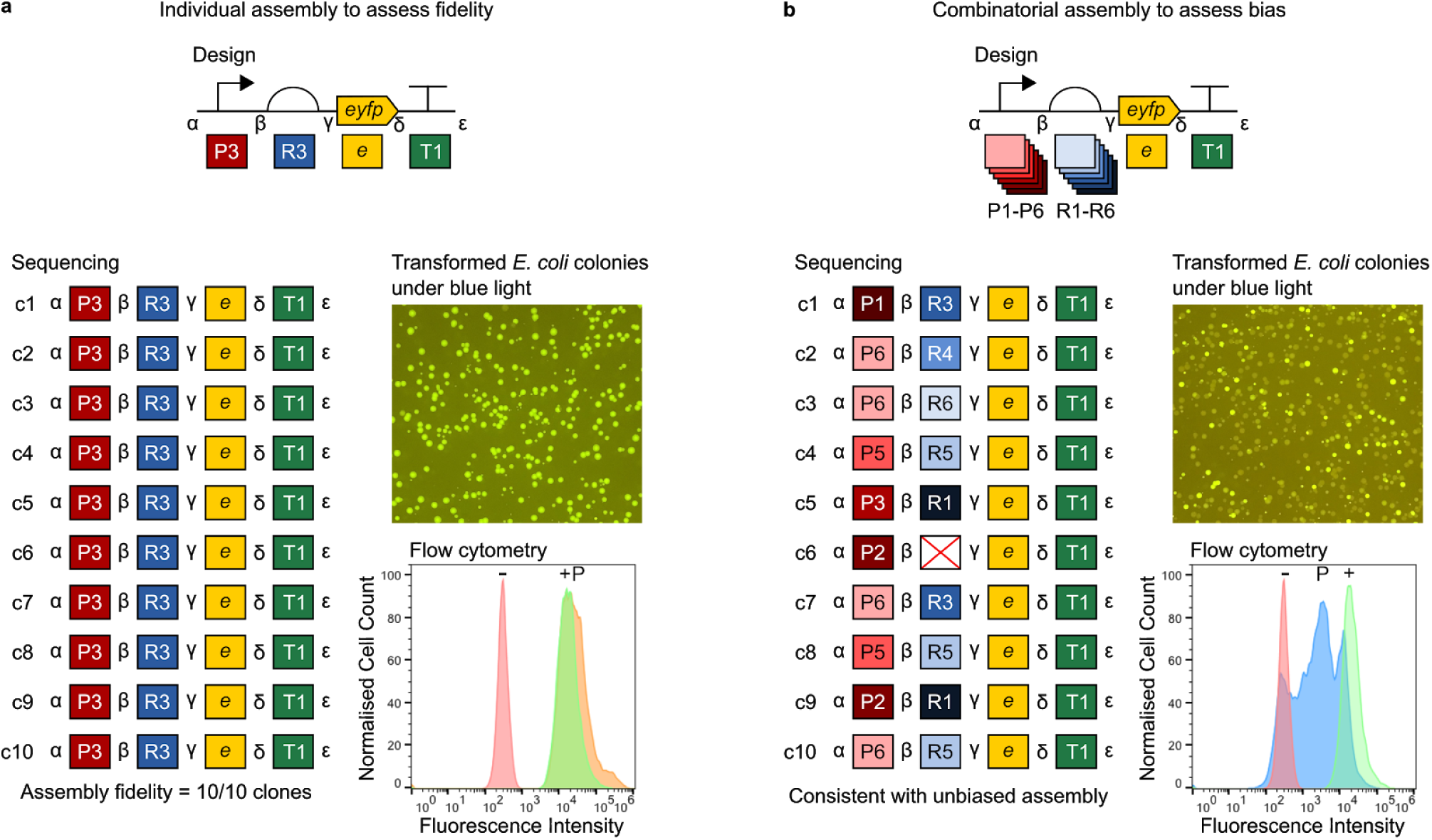
Assessment of fidelity and bias of multi-part Level 1 assembly. Constructs were assembled from Level 0 parts (in storage plasmids) into Level 1 vector pStA1AZ. Promoters, in descending order of strength: P1 = J23100, P2 = J23102, P3 = J23118, P4 = J23107, P5 = J23116, P6 = J23113. RBSs, in descending order of strength: R1 = RBSc44, R2 = RBSc33, R3 = RBSc13, R4 = RBSc58, R5 = RBSc42, R6 = RBSc36. CDS: e = *eyfp.* Terminator: T1 = L3S2P55. Representative images of *E. coli* DH10B colonies transformed with assembly reactions are shown illuminated by blue light to visualise fluorescence. In each experiment ten randomly-selected colonies (c1-c10) were analysed by sequencing (with primers oligoGT234 and oligoGT235) and flow cytometry. Sequencing results show expected intact parts and fusion sites. Flow cytometry histograms show fluorescence intensity of 10,000 events (cells) normalised to the maximum (in order to visualise distribution rather than absolute values) for wild-type *E. coli* DH10B as a negative reference (-), the most fluorescent of the ten clones as a positive reference (+) and a pool of several hundred transformants (P). **(a)** Assessment of assembly fidelity using assembly of an individual P3-R3-e-T1 expression unit. Two colonies were randomly selected for analysis from each of five replicate assemblies. The flow cytometry positive reference (+) strain is c3. **(b)** Assessment of assembly bias by combinatorial assembly of EYFP expression units using six promoters P1-P6, six RBSs R1-R6, *eyfp* and terminator T1. Ten colonies from one assembly were randomly selected for analysis. The flow cytometry positive reference (+) strain is c1. In the sequencing results the red cross indicates a misassembly between β and γ in place of the UTR/RBS.

For the experiment to assess fidelity, promoter P3, RBS R3, terminator T1 (Table S3) and the *eyfp* CDS were assembled into the Level 1 vector pStA1AZ (Table 1) using a one-pot assembly reaction including both SapI and ligase, as described in the Materials and Methods section. After incubation, the reaction mixture was used to transform *E. coli* DH10B. Five independent replicates of this experiment were performed. Blue/white screening using the *lacZ*α gene of pStA1AZ showed approximately one blue colony (containing the unmodified pStA1AZ vector) per 1000 white colonies. The fluorescence of colonies on agar plates was visualised by illuminating the plates with blue light, which showed a uniform level of fluorescence among colonies (Figure 2a). Two colonies from each of the five replicate experiments were selected at random for analysis. Sequencing of plasmids purified from these ten clones showed assembly of the expected parts and correct assembly of all five fusion sites in all ten constructs (Figure 2a). The fluorescence of cells of all ten clones, and a pool of several hundred transformant clones from one transformation, was determined by flow cytometry analysis of mid-exponential growth phase cultures. The fluorescence of cells in the pool was very similar to the fluorescence of cells in the ten verified, isolated clones (the histogram in Figure 2a shows one clone, the other clones are shown in Figure S21). These results indicate high fidelity of assembly of the Level 1 architecture using five 3 bp fusion sites in a one-pot reaction containing SapI and ligase.

For the experiment to assess bias, an equimolar mixture of all six promoters, an equimolar mixture of all six RBSs, CDS *eyfp* and terminator T1 were combinatorially assembled into the Level 1 vector pStA1AZ using the same procedures for assembly, transformation and analysis as described above for the fidelity experiment, except that the ten colonies selected at random for analysis were from a single assembly and transformation. Colonies on agar plates illuminated by blue light visibly showed diverse fluorescence (Figure 2b). Sequencing of plasmids purified from the ten clones showed assembly of expected parts and correct assembly of all five fusion sites in nine of the ten constructs. Construct c6 represents a misassembly, as the sequence indicates a spurious ligation of a fragment of *E. coli* genomic DNA and a fragment of vector DNA in place of the UTR/RBS part between the β and γ fusion sites (Figure 2b). The sequences of the nine correctly-assembled constructs show a variety of combinations of the possible promoters and RBSs. The fluorescence of cells in a pool of several hundred transformant clones, determined by flow cytometry as before, spanned a broad range of values from the negative control (pStA1AZ) to the positive control (which was the most fluorescent of the ten isolated clones in this experiment). These results are consistent with unbiased combinatorial assembly using the Level 1 architecture in a one-pot reaction.

### A streamlined assembly hierarchy to minimise the number of destination vectors required for new contexts and organisms

We set out to develop a multi-level hierarchical assembly framework incorporating the validated Level 1 architecture to allow assembly of constructs containing multiple expression units. The structure of the hierarchy is determined by the identity and arrangement of fusion sites within and between levels. In a Golden Gate-type multi-level assembly, vectors at each intermediate level (each level except the first or last) have both acceptor fusion sites for incorporation of parts from the preceding level, and donor fusion sites for subsequent excision of the assembled construct to be used as a part in the next assembly level. The Start-Stop Assembly Level 1 architecture (Figure 1a) requires the first (α) and last (ε) fusion sites to be the same in every case, which suits the assembly of individual expression units with defined, common, modular features. In contrast, most multi-level DNA assembly frameworks do not require a specific pair of fusion sites to be used as the first and last fusion sites in all assemblies at each level, but instead accommodate different combinations of first and last fusion sites, as a way of flexibly allowing different numbers of parts to be assembled together. This is achieved by providing sets of multiple alternative vectors at later levels, each with a different pair of acceptor fusion sites (for example, see Figure 3b). This approach is effective, but has a major drawback, as it means that these sets of multiple alternative vectors must be constructed for each different destination context, such as plasmids with different copy numbers, or shuttle vectors for different organisms, or different integration sites in an organism. This issue is significant for the use of multi-part DNA assembly in metabolic engineering, because there are many different organisms and contexts in which it would be useful to deploy assembled constructs encoding metabolic pathways. This issue does not appear to have been addressed in the design of previous modular multi-part assembly methods. We designed a different hierarchy to overcome this issue at the second multi-part assembly level, Level 2, where it would be most significant.

**Figure 3.**
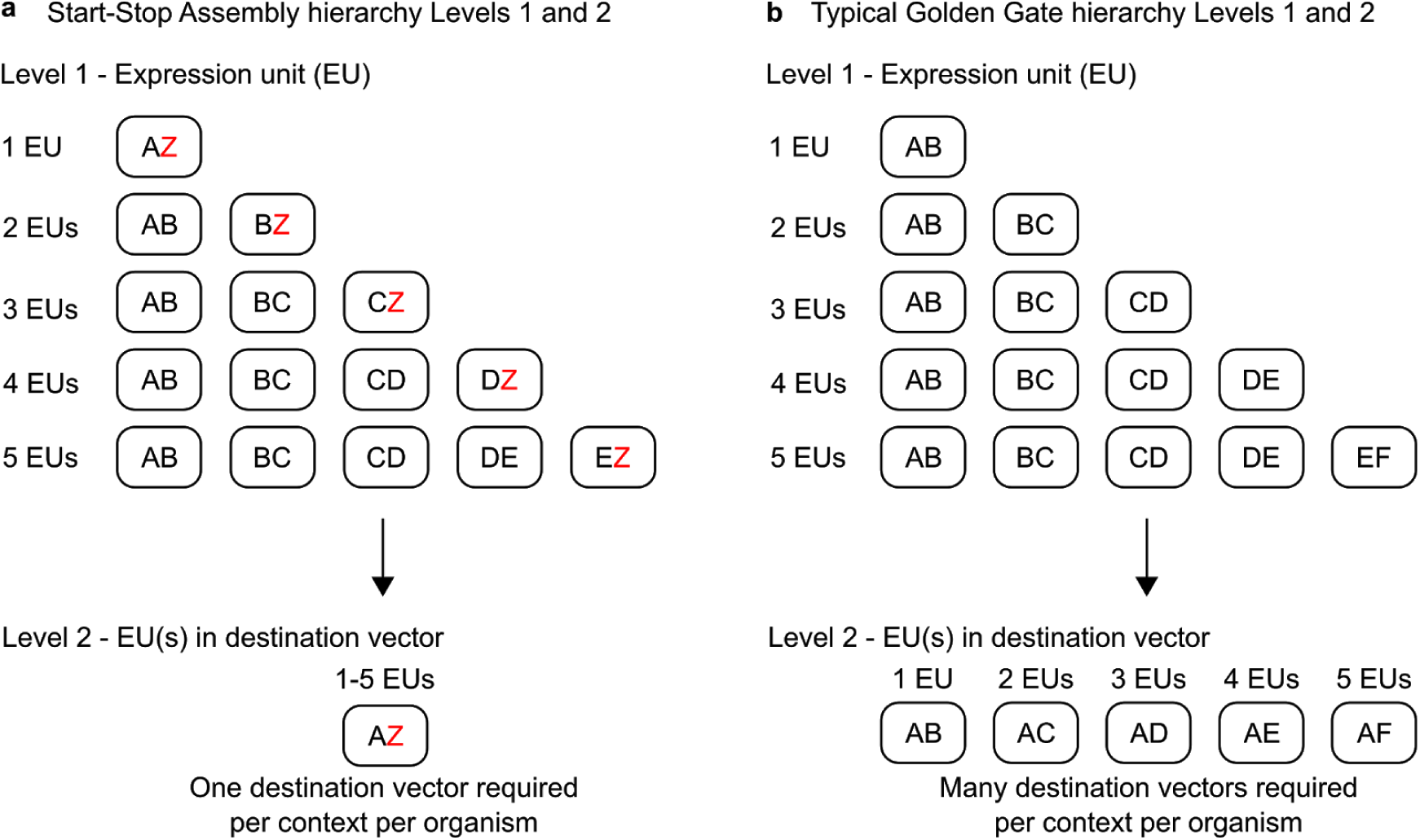
Alternative hierarchies achieved by alternative organisation of Level 1 donor and Level 2 acceptor fusion sites. Rounded rectangles represent vectors. Letters represent Level 1 donor fusion sites and Level 2 acceptor fusion sites, **(a)** Start-Stop Assembly uses alternative ‘Z vectors’ at Level 1 to minimise the number of destination vectors at Level 2. By assembling the last expression unit (of those being assembled at Level 1) in a Z vector at Level 1, in which the last donor fusion site is Z, then the recipient Level 2 vector can always use the same acceptor fusion sites, A and Z. While this requires more Level 1 vectors, it requires only one Level 2 vector. If alternative destination vectors are required for different organisms or contexts, only one new Level 2 vector needs to be constructed, **(b)** The typical Golden Gate framework requires multiple Level 2 vectors per context to accommodate all the possible Level 1 fusion site configurations. Therefore if an alternative destination vector is required several new vectors will need to be constructed per context or organism.

Level 2 is the destination (final) level for many assemblies. A typical Golden Gate hierarchy (Figure 3b) allowing assembly of up to five expression units at Level 2 would require five alternative combinations of acceptor fusion sites, and therefore five alternative destination vectors, per destination context. Instead, in Start-Stop Assembly we specified a hierarchy in which one particular pair of fusion sites, named A and Z, will always be the acceptor sites for Level 2 assembly of Level 1 parts into a Level 2 vector (Figure 3a). These are a subset of the donor fusion sites of Level 1 vectors (A, B, C, D, E and Z) the rest of which (B, C, D and E) are used for part-part junctions, but not for part-vector junctions. The purpose and advantage of this design choice is that only one vector (with A and Z acceptor fusion sites) is required at Level 2 regardless of the number of units being assembled (up to the five-unit maximum limit of Level 2, discussed later). Therefore any new destination context (such as a shuttle vector for a different organism) requires only one new Level 2 vector to be constructed (Figure 3a), instead of another set of five alternative Level 2 vectors (Figure 3b). For assembly into A and Z fusion sites at Level 2, the first Level 1 part to be assembled must begin with an A fusion site, and the last Level 1 part to be assembled must end with a Z fusion site. To achieve this while still allowing different numbers of expression units to be assembled together, one of the Level 1 vectors used in the assembly must be varied depending upon the number of units to be assembled. Specifically, for the last expression unit in a series of units to be assembled, the standard Level 1 vector is replaced by a ‘Z vector’, in which the second of the two fusion sites is replaced by a Z fusion site (Figure 3a). For example, for the third (and last) unit in a series of three units, the standard Level 1 vector, with C and D acceptor fusion sites, is replaced by an alternative Z vector, with C and Z acceptor fusion sites (Figure 3a, Figure 4, Figure S13).

**Figure 4.**
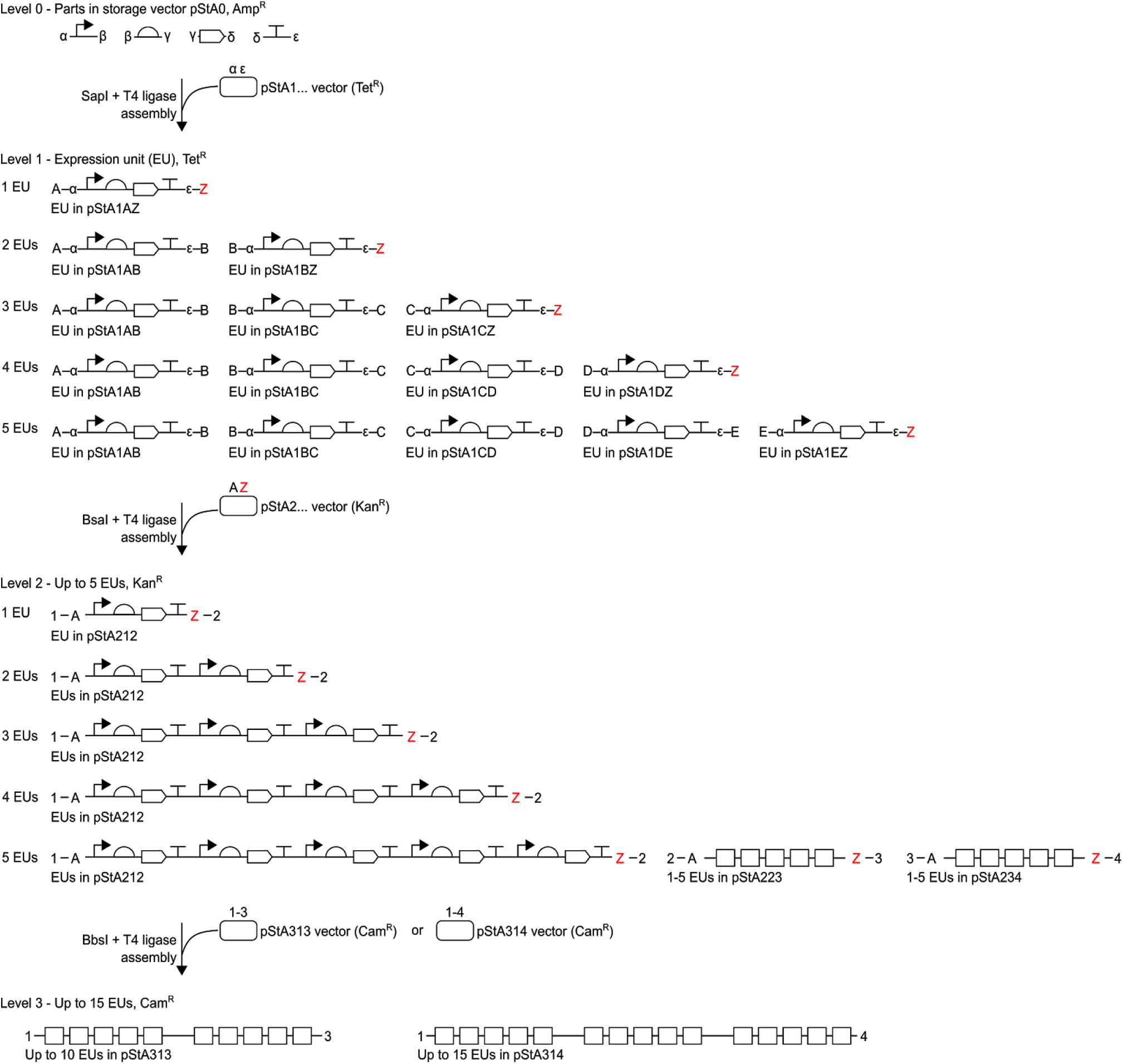
Detail of multi-part hierarchical assembly and fusion sites used in Start-Stop Assembly. Up to 15 expression units can be hierarchically assembled from basic parts (promoters, RBSs, CDSs and terminators) stored at Level 0. Level 1 expression units are assembled from four Level 0 parts, Level 2 constructs are assembled from up to five Level 1 expression units, and Level 3 constructs are assembled from up to three Level 2 constructs. Different type IIS restriction endonucleases (SapI, BsaI or BbsI) and different antibiotic-resistance markers (Amp^R^, Tet^R^, Kan^R^ or Cam^R^) are used for each assembly level as shown. Acceptor fusion sites are shown inside donor fusion sites at Level 1 and Level 2. Only donor fusion sites are shown at Level 0, and only acceptor fusion sites are shown at Level 3. As described in the text, alternative sets of vectors are used at Level 1 depending upon the number of expression units being assembled, such that the first and last Level 1 donor fusion sites are always A and Z, respectively, which are always the Level 2 acceptor fusion sites. Multiple Level 2 vectors (pStA212, pStA223, pStA234) with different donor fusion sites are only needed if the assembly will be continued to Level 3, in order to assemble more than five expression units. Alternative Level 3 vectors (pStA313, pStA314) with different acceptor fusion sites allow for assembly of two or three Level 2 constructs.

Effectively, the streamlined hierarchy moves the requirement for additional alternative vectors from a destination level (Level 2) where it is problematic as described above, to an intermediate, non-destination level (Level 1) where it is not problematic because the core set of vectors (including alternative Z vectors) we provide here (Table 1) can be used for assembly of various numbers of units regardless of any changes to the final assembly destination vector. The user need only construct a single new (Level 2) destination vector per new context or organism.

Level 2 was limited to assembly of a maximum of five Level 1 parts (expression units) in order to maintain a high assembly efficiency, which is important to ensure that libraries containing large numbers of clones are readily obtained by combinatorial assembly. To accommodate cases which require assembly of greater numbers of expression units, we extended the hierarchy to Level 3, which supports the assembly of up to 15 expression units (Figure 4). Level 2 vectors therefore contain donor fusion sites (referred to by numbers 1, 2, 3 or 4) to allow excision of Level 2 assembled constructs for use as parts in Level 3 assembly, and Level 3 vectors contain different combinations of the corresponding acceptor fusion sites. Extending the hierarchy to Level 3 requires additional Level 2 vectors (Figure 4, Table 1). However, the advantage of the streamlined hierarchy described above (and emphasised in Figure 3) is not lost, because when Level 3 assembly is used, the destination vector will be at Level 3, and Level 2 will serve as an intermediate level, not a destination level, so the Level 2 vectors already provided here can be used. Therefore it is still only necessary for the user to construct one or possibly two new (Level 3) destination vector(s) per new context or organism.

### Implementation of the complete multi-level Start-Stop Assembly system

Having established the details of the architecture of Level 0 and Level 1 which provide for functionally-scarless assembly of each expression unit (Figure 1a), and the overall assembly hierarchy for all levels (Figure 1b, Figure 4), we proceeded to resolve all remaining design details and implement Start-Stop Assembly as a complete modular DNA assembly system with a widely-applicable set of 15 core vectors (Figure 1b, Figure 4).

First, we defined the details of Level 2 assembly and Level 3 assembly. Unlike Level 1 assembly of expression units from Level 0 parts, fusion sites for Level 2 assembly and Level 3 assembly can be located outside regions that are acutely sensitive to scar sequences. Therefore for Level 2 and Level 3 it was not necessary to apply detailed design constraints like those described earlier for the Level 1 architecture and assembly. Instead, we chose type IIS restriction endonucleases and fusion site sequences for Level 2 assembly and Level 3 assembly from among those reported and validated previously. We chose the type IIS enzyme BsaI for Level 2 assembly and BbsI for Level 3 assembly, both of which are commonly used in Golden Gate-based methods (4, 59). Restriction of DNA by BsaI or BbsI yields cohesive DNA ends with 4 bp overhangs, which correspond to 4 bp fusion sites. We assigned 4 bp sequences to the fusion sites A (GGAG), B (AATG), C (AGGT), D (GCTT), E (CGCT) and Z (TACT) for Level 2 assembly of Level 1 parts into a Level 2 vector. We assigned 4 bp sequences to the fusion sites 1 (TGCC), 2 (ACTA), 3 (TTAC) and 4 (CGAG) for Level 3 assembly of Level 2 parts into a Level 3 vector. All these sequences have previously been shown to provide high-fidelity assembly (4).

Each of the 15 core Start-Stop Assembly vectors contain a similar assembly cassette to facilitate the initial cloning of parts at Level 0 and their subsequent hierarchical assembly through one or more further Levels (Table 1, Figure 4). In the centre of each cassette is a *lacZα* gene flanked by the relevant acceptor fusion sites, with outward-facing recognition sites for the corresponding type IIS restriction endonuclease, positioned appropriately to ensure cohesive DNA ends correspond to the fusion sites. This arrangement allows blue/white screening (in suitable alpha-complementing *E. coli* strains) for replacement of the *lacZα* gene upon successful assembly using the acceptor fusion sites. Outside the acceptor fusion sites are the relevant donor fusion sites, with inward-facing recognition sites for the corresponding type IIS restriction endonuclease (except in Level 3 vectors, which lack donor fusion sites) positioned appropriately to ensure cohesive DNA ends correspond to the fusion sites. Finally, the fusion sites are flanked by the strong, *rho*-independent transcriptional terminators T_0_ of phage lambda and T_1_ of the *rrnB* operon of *E. coli* (60). These terminators transcriptionally insulate cloned parts and assembled constructs from the rest of the vector, preventing transcriptional ‘read-in’ and/or ‘read-out’ which might interfere with the function of either assembled sequences or the vector.

We constructed a core set of 15 vectors including assembly cassettes, with fusion sites described above, in combinations shown in Table 1 and Figure 4. To avoid carry-over of plasmids from one level to the next during assembly, we designed each level to encode a different antibiotic resistance from the next, as in other multi-level assembly systems (4, 5, 59). The Level 0 vector encodes ampicillin resistance (Amp^R^), Level 1 vectors encode tetracycline resistance (Tet^R^), Level 2 vectors encode kanamycin resistance (Kan^R^) and Level 3 vectors encode chloramphenicol resistance (Cam^R^). We designed Level 1, Level 2 and Level 3 vectors to use the low copy-number replicon p15A (10-12 copies per cell (61)) in order to minimise the burden imposed on cells by the complete, functional expression units assembled at these levels. Assembled constructs encoding metabolic pathways could cause such burden directly through protein over-expression, and/or indirectly through diversion of metabolite flux into the expressed pathway. The use of low copy-number replicons to reduce these deleterious effects should improve the success of assembly of constructs expressing metabolic pathways and minimise bias in combinatorial assembly. Level 0 (part storage) does not involve functional expression units, so a low-copy number replicon would be less useful at Level 0. Instead, to allow more convenient preparation and sequence-verification of parts cloned in the Level 0 vector, a high copy-number replicon pMB1 (500-700 copies per cell) was used. These combinations of antibiotic resistances and replicons were achieved by constructing the 15 core vectors from the classic vectors pUC19, pACYC177 or pACYC184 (Supplementary Materials and Methods). SapI, BsaI and BbsI restriction sites are reserved for assembly, so were removed from all the vectors by conventional methods (Supplementary Materials and Methods), except for the intended positions in the assembly cassettes. Validated primers for sequencing parts or assembled constructs at all levels are listed in Table S4. In order to apply Start-Stop Assembly to new destination contexts (such as organisms or integration sites), alternative Level 2 or Level 3 destination vectors can be constructed simply by installing suitable combinations of restriction sites and fusion sites (typically by transferring the entire assembly cassette from an existing Level 2 or Level 3 vector; Figure S6) into the alternative vector backbone, and removing any other BsaI and/or BbsI sites.

Start-Stop Assembly allows assembly of independent monocistronic expression units, which have advantages over operons, particularly as they allow greater independence of expression units, and more fine-grained control of expression, as both promoters and RBSs can be varied for each unit. However, there are scenarios in which operon configurations may be preferred, such as the co-regulation of multiple expression units by a single regulated promoter, as found in natural operon systems (62, 63), or to cause the expression levels of multiple units to be partially linked in combinatorial designs (30–32), or to allow a metabolic pathway or other system to be optimised under the control of one promoter and then subsequently, once optimised, placed under the control of a different promoter suited to a particular application. To allow operon configurations within the same overall framework, we generated short spacer sequences which can be used in place of a promoter and/or terminator in Level 1 assembly of an expression unit. This approach can be used to generate Level 1 units suitable for the start of an operon (with a spacer in place of a terminator), the end of an operon (with a spacer in place of a promoter), or the middle of an operon (with spacers in place of both promoter and terminator). A set of 16 spacers designed to be orthogonal and biologically-neutral, each 12 bp in length, were generated using the R2oDNA designer software (64) (Table S5). As the spacers are short, it is convenient to implement these spacer parts as linkers, by annealing partially complementary pairs of oligonucleotides to obtain double-stranded linkers with cohesive ends corresponding to fusion sites (Table S5). For each spacer sequence, two such linkers were designed, the first with α and β cohesive ends, suitable for Level 1 assembly in place of a promoter; and the second with δ and ε cohesive ends, suitable for Level 1 assembly in place of a terminator. These linkers can be used directly in a Level 1 assembly reaction (Supplementary Materials and Methods).

### Combinatorial Start-Stop Assembly of metabolic pathways provides exploration of design space, phenotypic diversity and high-performance pathway configurations

We set out to establish Start-Stop Assembly as an effective and generally-applicable platform for the implementation and optimisation of metabolic pathways. We aimed to validate (i) combinatorial assembly of constructs encoding (ii) effective metabolic pathways (iii) composed of numerous enzymes; (iv) in monocistronic, operon and hybrid configurations; and (v) with wide exploration of design space demonstrated by observable phenotypic properties. Metabolic pathways for the biosynthesis of carotenoids are ideal for these purposes, because differently-coloured compounds are formed at several steps in the pathways, including the final products. These red, yellow and orange colours effectively serve as an indirect, semi-quantitative readout of the balance between the fluxes in the pathway, caused by the relative expression levels of the enzymes catalysing the various steps. Diversity and performance of carotenoid pathway configurations is therefore readily observed. Carotenoid pathways have been used in this way previously (32, 40, 65). Furthermore, carotenoids are valuable compounds with useful properties used in the pharmaceutical, food and cosmetic industries (66).

We used Start-Stop Assembly to combinatorially assemble, in *E. coli*, five libraries of constructs encoding carotenoid metabolic pathways. Four of these libraries (pGT531-534, Figure S16-19) encode pathways for the product β-carotene (Figure S15a) using four heterologous CDSs (*crtE*, *crtB*, *crtI* and *icyB*) plus the native *E. coli* CDS *dxs* (encoding an enzyme in the upstream MEV pathway which is known to be rate-limiting in *E. coli* (32, 67-69)) making five CDSs in total. The four β-carotene pathway libraries use different monocistronic, operon or hybrid configurations to confirm that these configurations perform as expected in Start-Stop Assembly. The fifth pathway library (pGT535) includes the same five CDSs as the β-carotene pathway libraries plus three additional CDSs (*idi*, *crtZ* and *crtW*), which extend the pathway to the product astaxanthin (Figure 5a), making eight CDSs in total (Figure S20). The eight CDSs (detailed in the Materials and Methods section) were cloned into the Level 0 vector pStA0 (Table S3). Pathway libraries were combinatorially assembled using these CDSs and the sets of six promoters, six RBSs and (individually) four terminators generated earlier (Table S3) in order to vary the expression of each enzyme widely. Combinatorial variation was introduced by using equimolar mixtures of the six promoter parts or six RBS parts at each appropriate position in Level 1 assemblies, then propagated to subsequent assembly levels through hierarchical assembly. In operon designs, spacers (described earlier, and shown in Table S5) were used in place of promoters and terminators at appropriate positions as shown in Figures S17-20, which also show the vectors used. The first β-carotene pathway library (pGT531) was assembled in a monocistronic configuration in the destination vector pStA212 (Figure S16). Due to the mixtures of six promoter parts and mixtures of six RBS parts used, the total combinatorial design space (the maximum possible library size) was 6^10^ (which equals 6.05×10^7^), which is the product of multiplying the number of parts (six) in the mixtures in each of ten combinatorial positions (Figure S16). In the second β-carotene pathway library (pGT532) the same five CDSs were used, but in an operon configuration, assembled in pStA223, accordingly with a much smaller total combinatorial design space of 6^6^ (which equals 4.67×10^4^, Figure S17). The third β-carotene pathway library (pGT533) was constructed in pStA234 in a hybrid configuration, composed of a three-unit operon and two monocistronic units, with a total combinatorial design space of 6^8^ (which equals 1.68×10^6^, Figure S18). The fourth β-carotene pathway library (pGT534) was also assembled in a hybrid configuration, composed of a four-unit operon and one monocistronic unit, which was assembled in pStA313 with a total combinatorial design space of 6^7^ (which equals 2.80×10^5^, Figure S19). The astaxanthin pathway library (pGT535) was combinatorially assembled in pStA314 in a hybrid configuration, composed of a four-unit operon, a monocistronic unit, and a three-unit operon. For the astaxanthin pathway library, the total combinatorial design space was 6^11^ (which equals 3.63×10^8^, Figure S20).

**Figure 5.**
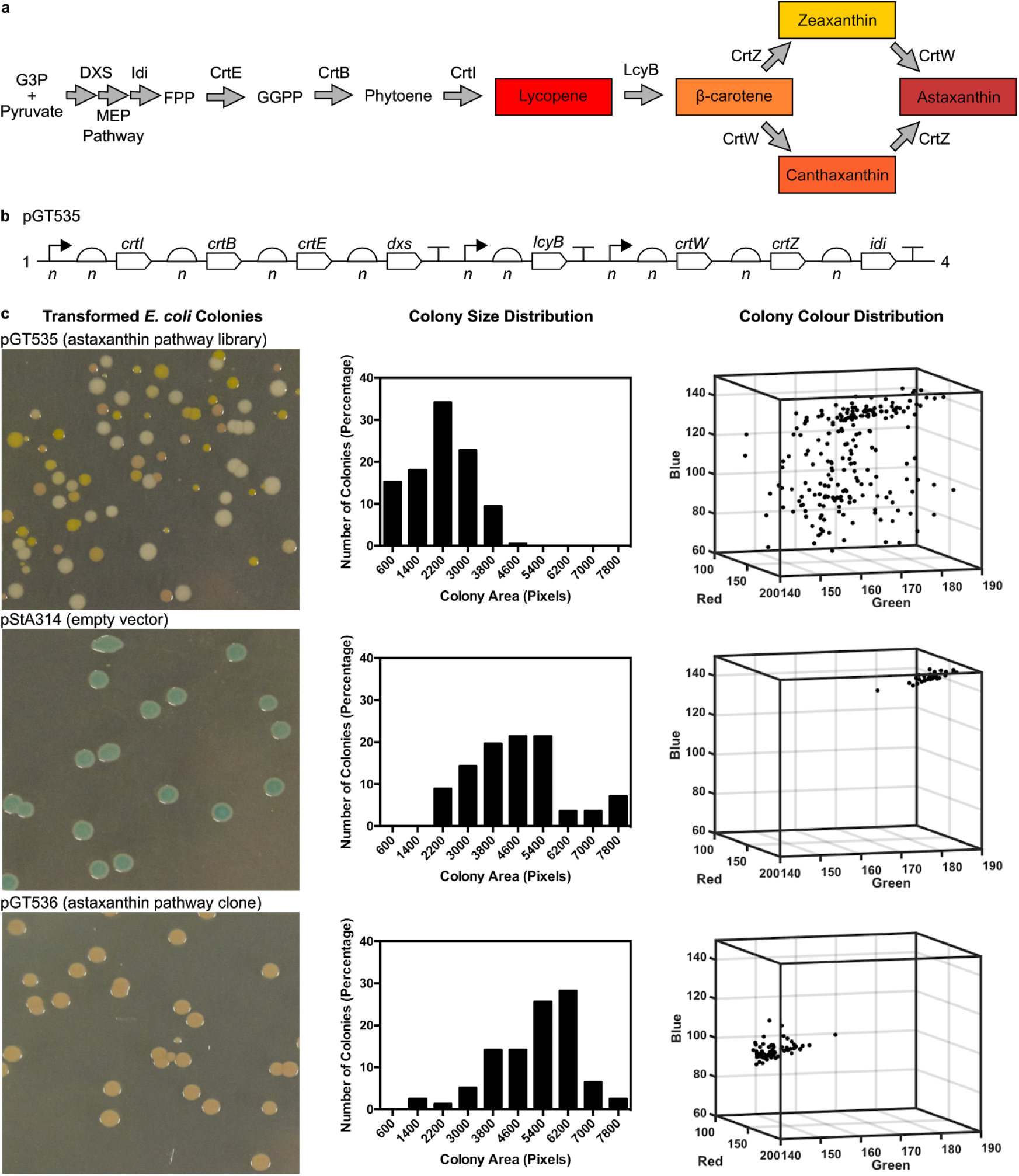
Combinatorial assembly of astaxanthin pathway using Start-Stop Assembly. Astaxanthin pathway showing the endogenous enzymes of the *E. coli* MEP pathway and heterologous enzymes. Coloured products are shown in boxes, enzymes included in the combinatorial assembly are shown above arrows. Abbreviations: G3P (glyceraldehyde 3-phosphate), FPP (Farnesyl pyrophosphate), GGPP (Geranylgeranyl pyrophosphate), **(b)** Design of astaxanthin pathway library pGT535, composed of eight expression units combinatorially and hierarchically assembled into Level 3 vector pStA314, as outlined in the text and shown in detail in Figure S20. Mixtures of six parts of varying strengths were used at each promoter and RBS position shown, as described in the text. These are shown as *n* in the figure to reflect the sample of the combinatorial design space present after hierarchical assembly, **(c)** Phenotypic variation among *E. coli* clones of the astaxanthin pathway library (pGT535) was compared to controls pStA314 (empty vector) and an isolated clone pGT536 from the astaxanthin pathway library pGT535. Phenotypic variation is shown using representative images of colonies, histograms of colony size (measured as colony area in pixels, x-axis values represent the upper limit of each histogram bin) and the distributions of colony colours (represented using red, green and blue values extracted from colony images).

In each case, the pathway libraries were generated by hierarchical assembly using one-pot assembly reactions at each step, then pooling all white colonies (typically 99% of colonies were white and 1% were blue) obtained at each intermediate step to prepare plasmid DNA for use as the combinatorial mixture of parts for the subsequent step. Each transformation yielded at least thousands of colonies, and typically tens of thousands of colonies, providing many-fold coverage at the Level 1 assembly steps (at which the maximum design space was 6^2^ = 36) and typically partial coverage at the Level 2 and Level 3 assembly steps (at which the design space was between 6^6^ (which equals 4.67×10^4^) and 6^11^ (which equals 3.63×10^8^), so the libraries of clones obtained as colonies on plates at the end of each hierarchical assembly represent a sample of the possible design space.

As the combinatorial libraries were designed to vary the expression of each enzyme in the carotenoid pathways widely, we expected to observe colonies of a range of colours and intensities of those colours caused by differing amounts of the various carotenoids. The astaxanthin pathway library in particular could produce up to five coloured compounds (Figure 5a, 5b). We also expected colony sizes to vary widely, reflecting differing growth impairments caused by protein over-expression and/or by central metabolites being directed away from native metabolism to carotenoid biosynthesis. Diversity of colony colour and size were both clearly evident by eye, and we analysed images of plates of colonies to measure these effects more objectively (Figure 5c, Figure S15). In comparison to controls (the empty vector pStA314 and an isolated clone pGT536 from the astaxanthin pathway library pGT535) the distribution of colony sizes in the astaxanthin pathway library pGT535 was shifted towards much smaller colonies, and the colonies showed a wide distribution of colours not seen in the controls (Figure 5c). Interestingly, colony colours were not associated with with colony size (linear regression *R*^2^ values <0.2, Figure S22), and colonies of similar colours but very different sizes were observed. Similar results were obtained with the four different β-carotene pathway libraries, among which some differences were observed in both colony size distributions and colony colour distributions (Figure S15 and S23), presumably caused by the different designs of the pathway libraries. Finally, as a simple measure of the effectiveness of the pathway variants obtained, and to place the observed variation in the context of previously-reported carotenoid biosynthesis, ten clones were picked at random from the astaxanthin pathway library and analysed for their production of lycopene, the first coloured carotenoid in the pathway (Figure 6). In simple, small-scale batch cultures over 24 h we observed different lycopene concentrations from 0 mg/g DCW to 108 mg/g DCW, a similar range to previous comparable studies (40, 70, 71), indicating that the variants obtained here represent effective pathway implementations.

**Figure 6.**
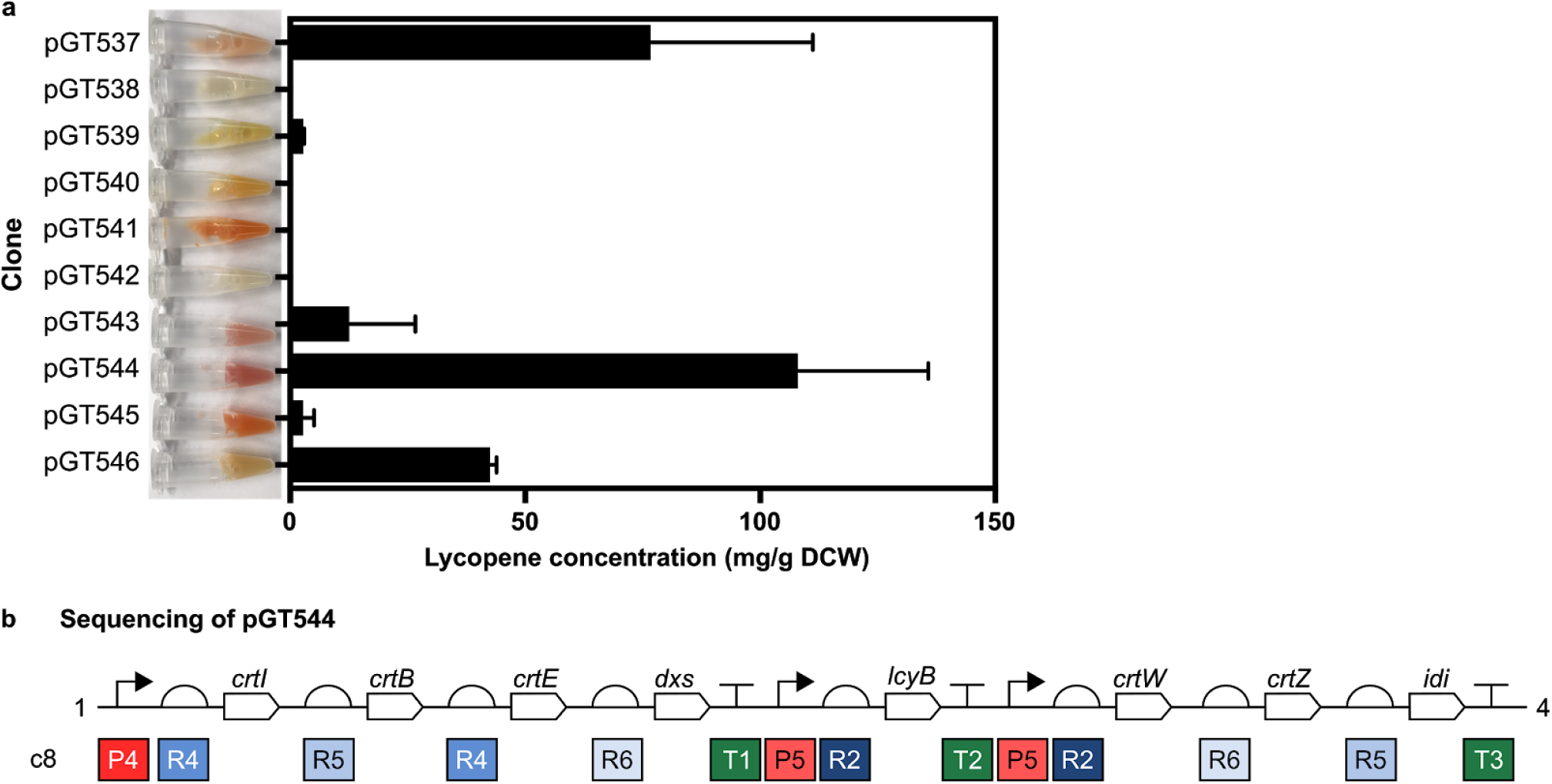
Lycopene content of *E. coli* clones from the astaxanthin library. **(a)** Ten *E. coli* colonies were randomly selected from the combinatorially-assembled astaxanthin pathway library pGT535 (Figure 5, Figure S20). Each clone was grown for 24 h in LB broth before lycopene content was determined as mg lycopene per dry cell weight (DCW). Error bars represent the standard deviation of three independent biological replicates. A photograph of a sample of the culture of each clone was taken (shown to left of graph), which shows the visible variation of colour in the ten cultures, attributed to their different profiles of the carotenoids produced in the astaxanthin pathway, **(b)** Sequencing data for the clone containing pGT544. Promoters, in descending order of strength: P4 = J23107, P5 = J23116. RBSs, in descending order of strength: R2 = RBSc33, R4 = RBSc58, R5 = RBSc42, R6 = RBSc36. Terminator: T1 = L3S2P55, T2 = L3S2P21, T3 = ECK120033737.

## DISCUSSION

All DNA assembly methods impose constraints of one form or another on the design and assembly of sequences and libraries, particularly the scars of modular assembly methods, and the low efficiency, bias and issues with repetitive sequences of bespoke assembly methods. Operating within the relevant constraints is a necessary compromise to use an assembly method and benefit from its advantages. However, is it generally preferable to avoid unnecessary design constraints where possible. Here, we develop and validate Start-Stop Assembly, which provides the same key advantages as other Golden Gate-type modular assembly frameworks; particularly efficient, unbiased, multi-part, hierarchical assembly; but also allows designs to be scarless at CDS boundaries, which are crucial, highly-sensitive sites where scars affect mRNA structure and the activity of the RBS (as demonstrated by our direct comparison of the impact of scars in Figure S1) and potentially other functional RNA features. By comparison to Start-Stop Assembly, the scars at CDS boundaries in other modular DNA assembly methods represent avoidable design constraints. The second key distinctive advantage of Start-Stop Assembly is the streamlined hierarchy, which means that typically only one new vector is required in order to assemble constructs for any new destination context (Figure 3). This should facilitate more rapid and convenient development of engineered metabolic pathways for diverse non-model organisms in order to exploit their industrial potential.

The main disadvantage of using start and stop codons for assembly is the incompatibility of this approach with the construction of fusion proteins from domains and/or tags, as this requires fusion sites which do not start or stop translation. Fortunately combinatorial construction of fusion proteins is infrequently used in most metabolic engineering. If required, individual fusion proteins could be included in Start-Stop Assembly simply by generating Level 0 CDS parts encoding complete fusion proteins in advance of assembly.

To the best of our knowledge, there is only one previous description of a DNA assembly method which is both modular and scarless, called ‘scarless stitching’ (31). This method uses Golden Gate-like restriction-ligation and fusion sites, but is necessarily limited to assembly of only two parts in each step due to an additional blunt restriction-ligation reaction used to delete ‘bridging’ (scar) sequences. Hierarchical assembly of metabolic pathways using scarless stitching would require many more steps in series and in parallel than multi-part methods like Start-Stop Assembly.

This work has focused on construction of prokaryotic expression constructs, but the system can readily be applied to construction of eukaryotic expression constructs using the same framework of fusion sites, hierarchy, storage vector and assembly vectors, simply by cloning suitable parts in the Start-Stop Assembly format and constructing a suitable destination vector.

The assembly and analysis of the carotenoid pathway libraries validates Start-Stop Assembly as a platform for the implementation and optimisation of metabolic pathways, as (i) the diversity of colony colour and colony size phenotypes reflects the intended wide exploration of design space, (ii) the effectiveness of pathway variants, reflected by carotenoid colours and lycopene concentrations, indicates the absence of emergent issues caused by novel combinations and arrangements of fusion site sequences, and (iii) all 15 core assembly vectors (Table 1) were successfully used during the construction of these five pathway libraries. The aim of these single-round experiments was to validate the approach, not to maximise production, although the results are encouraging and compare well to other reports (40, 70, 71). The observed lack of association between colony colour and colony size (Figures S22 and S23) suggests that different variants encode similarly productive pathways while imposing very different growth impairments on the host cell, and may reflect trade-offs between transcription and translation (37, 72). This indicates that optimisation of metabolic pathways through their enzyme expression profile is important to obtain high-performing, low-burden pathway implementations, and is readily achieved by combinatorial assembly of suitable parts.

Multi-part DNA assembly methods are relatively complex by comparison to conventional restriction cloning, which poses a barrier to their widespread adoption, particularly if the advantages are not clear and the constraints are onerous, or appear so. Thus in this report we have aimed to provide sufficient detail to make Start-Stop Assembly very accessible, including to those who are unfamiliar with other multi-part DNA assembly methods. Much of this detail is found in the Supplementary Materials which include numerous figures and tables, a quick-start guide, and a laboratory protocol. The assembly of carotenoid pathway libraries in this study provides an exemplar and template for those seeking to use combinatorial assembly to construct and optimise metabolic pathways. We make available to the community (through Addgene) a kit of the 15 core storage and assembly vectors (Table 1) as well as all the promoters, RBSs, CDSs and terminators (in storage plasmids, Table S3) used successfully in the examples shown here. We suggest that a straightforward way for a research group to establish combinatorial multi-part assembly of metabolic pathways is to first directly reproduce one of the carotenoid libraries described here as a positive control, which provides immediate visual feedback of success in the form of colony colours and sizes, and then to simply repeat the assembly replacing carotenoid pathway CDS parts with CDS parts for the pathway of interest.

As combinatorial multi-part assembly becomes more routine and widespread, supporting the construction of large libraries, the challenge and focus of effort in the development of metabolic pathways is changing. Assembly itself is straightforward, but even the largest libraries inevitably represent only small samples of the very large possible design spaces, so it is not feasible to exhaustively screen all possible variants for the best-performing and least burdensome pathway variants. Effective strategies are needed first to design the granularity and distribution of fractional sampling of large pathway design spaces given a *priori* information, and then to iteratively optimise designs by moving through design space towards progressively better pathway variants. Such strategies have become the main subject of various recent studies (31, 73–75). This is a significant development reflecting the evolving role of systematic synthetic biology approaches in underpinning increasingly effective metabolic engineering and industrial biotechnology.

## ASSOCIATED CONTENT

Supplementary Materials are available: Figures S1-23, Notes S1-3 and Tables S1-8. We suggest that a reader intending to use Start-Stop Assembly first refers to Note S1, which provides a quick-start guide.

## AUTHOR INFORMATION

### Corresponding Author

John T. Heap

### Author Contributions

GT, PM and JH designed the study; GT performed experiments; GT and JH prepared the manuscript with input from PM.

### Notes

The authors declare no competing financial interest.

## ACKNOWLEDGEMENTS

The authors thank Dr. Soo Mei Chee for LC-UV analysis and thank Dr. Ciarán Kelly, Dr. Linda Dekker, Lara Sellés Vidal and Dr. Alexandra Faulds-Pain for useful discussions and critical input.

